# Imprecise recombinant viruses evolve via a fitness-driven, iterative process of polymerase template-switching events

**DOI:** 10.1101/2021.06.01.446546

**Authors:** Kirsten Bentley, Fadi G. Alnaji, Luke Woodford, Siân Jones, Andrew Woodman, David Evans

## Abstract

Recombination is a common feature of many positive-strand RNA viruses, playing an important role in virus evolution. However, to date, there is limited understanding of the mechanisms behind the process. Utilising *in vitro* assays, we have previously shown that the template-switching event of recombination is a random and ubiquitous process that often leads to recombinant viruses with imprecise genomes containing sequence duplications. Subsequently, a process termed resolution, that has yet to be mechanistically studied, removes these duplicated sequences resulting in a virus population of wild type length genomes. Using defined imprecise recombinant viruses together with Oxford Nanopore and Illumina high throughput next generation sequencing technologies we have investigated the process of resolution. We show that genome resolution involves subsequent rounds of template-switching recombination with viral fitness resulting in the survival of a small subset of recombinant genomes. This alters our previously held understanding that recombination and resolution are independent steps of the process, and instead demonstrates that viruses undergo frequent and continuous recombination events over a prolonged period until the fittest viruses, predominantly those with wild type length genomes, dominate the population.

**Author Summary:** Viruses with positive-sense RNA genomes, such as poliovirus, have several mechanisms by which they evolve. One of these is the process of recombination involving the large-scale exchange of genetic information. Recombination occurs during replication when the viral polymerase switches from copying one genome to another. However, the polymerase does not always accurately switch between the two, resulting in sequence duplications or deletions, and genomes that are referred to as imprecise. Over multiple rounds of replication sequence duplications are lost and genomes are resolved to wild type length, but it is unclear how this occurs. Here we used synthetic polioviruses containing defined sequence duplications to determine that the genome population undergoes repeated rounds of recombination until sequence duplications are lost and viruses with precise, wild type length genomes are selected for. This selection is based on the overall fitness of the virus population, with less fit imprecise viruses evolving more quickly. Our study suggests that recombination is a continual process where virus fitness drives the selection of a small subset of recombinant variants. These data are important for understanding how novel viruses evolve via recombination and how this process can be blocked to prevent novel and dangerous pathogens from arising.

## Introduction

RNA viruses usually exist as quasispecies due to error-prone RNA-dependent RNA polymerases (RdRps) which misincorporate nucleotides during genome replication. In many RNA viruses, the process of recombination generates additional diversity within the virus population. Recombination allows for a larger exchange of genetic material between closely related viruses and is proposed to be a mechanism by which genomes can be rescued from error catastrophe, which would otherwise have resulted from the accumulation of deleterious mutations introduced by the RdRp [1–3]. Recombination is widely accepted to occur via a copy-choice mechanism involving the RdRp switching from one template (the donor) to another (the acceptor) during negative-strand synthesis [4]. Sequence identity and RNA structure have been proposed to influence the template-switching event, although evidence is often contradictory and may reflect analysis of highly selected populations (reviewed in [5]).

Poliovirus, the prototype species C enterovirus of the *Picornaviridae* family, has proven to be a useful tool for the study of recombination. It has long been known that the three serotypes frequently recombine [6, 7] and there are well-established reverse genetic techniques allowing markers for recombination to be engineered into the genome [8, 9]. Poliovirus, like all enteroviruses, has a positive-strand RNA genome of approximately 7.5 Kb. The genome encodes a single polyprotein flanked by 5′ and 3′ untranslated regions (UTR) containing the signals for translation and replication. The polyprotein is co- and post-translationally cleaved to yield the structural proteins (P1: VP4, VP3, VP2, VP1) and the non-structural proteins (P2: 2A^pro^, 2B, 2C and P3: 3A, 3B^Vpg^, 3C^pro^ and 3D^pol^).

We have developed an *in vitro* recombination assay – the poliovirus CRE-REP assay – that solely generates recombinant progeny viruses [10]. The CRE-REP assay requires co-transfection of two RNA molecules that independently cannot generate viable virus progeny; (1) a poliovirus serotype 1 (PV1)-based replicon in which the structural proteins have been replaced with a luciferase marker gene, and (2) a full-length poliovirus serotype 3 (PV3) genome in which the 2C *cis*-acting replication element (CRE; [11]) has been mutated so that the virus can no longer synthesise new positive-strand genomes [12]. When co-transfected into permissive cells the two non-infectious genomes produce negative strands, which can recombine to yield a population of viable recombinant viruses. Using this assay, we postulated that recombination is a biphasic process [10]. First, an RdRp strand-transfer event produces genomes that may contain sequence duplications of up to several hundred nucleotides. The imprecise and apparently random nature of this recombination event argues strongly against sequence identity or structure being a major influence [13]. Since *in vivo* recombination results in genome-length products we proposed that strand transfer is followed by a second event we termed resolution, in which sequence duplications are lost via an iterative process. This could be demonstrated experimentally *in vitro* [10]. Resolution results in a virus population containing a variety of precise junctions between donor and acceptor templates [10]. Mechanistically the process of resolution is poorly defined, but two obvious scenarios can be envisaged: an *in-cis* internal deletion event involving individual genomes or an *in-trans* process requiring two or more genomes (Fig 1).

**Fig 1.**
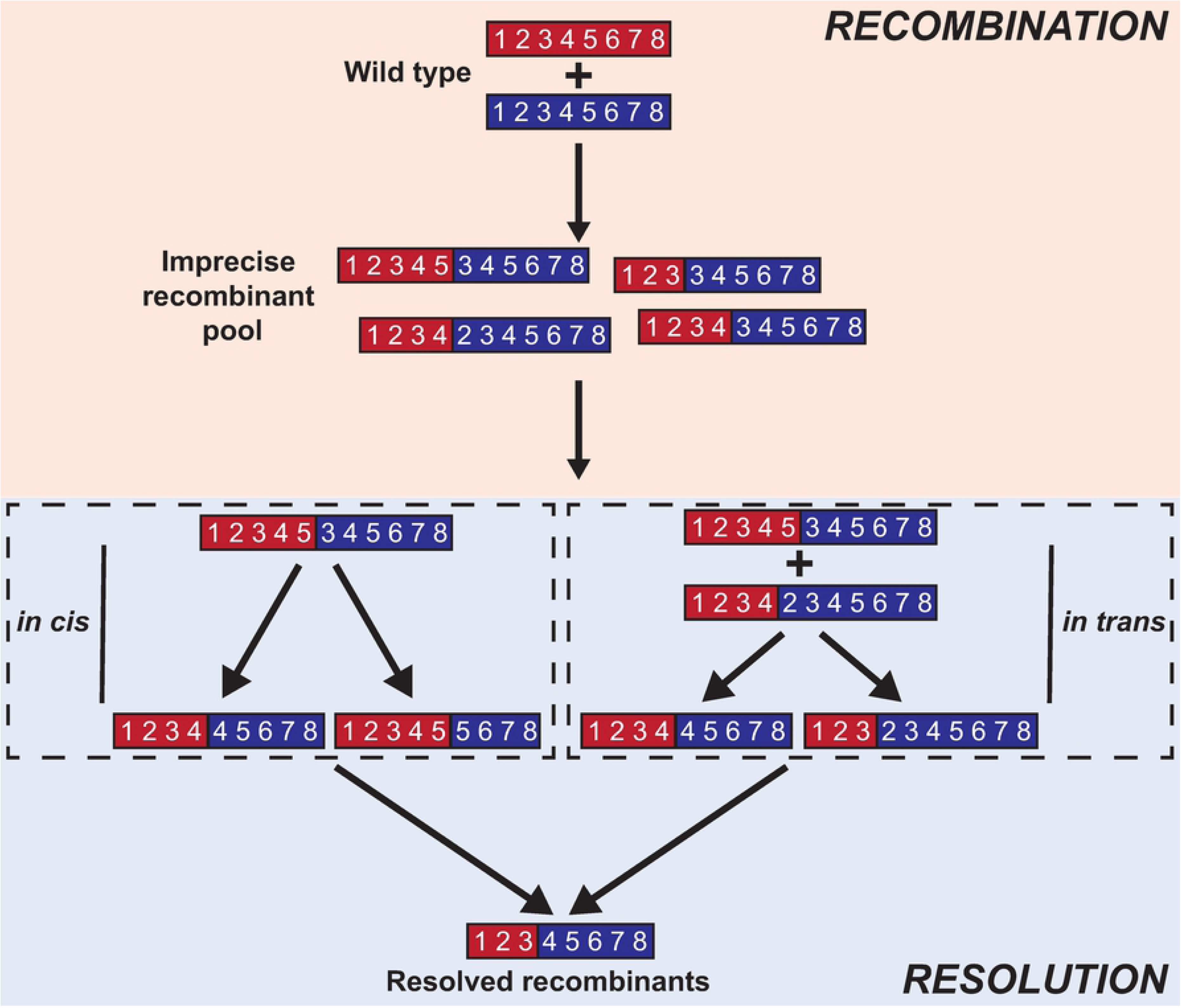
Model of recombination and resolution. Following a recombination (strand transfer) event a mixed population of viruses is generated containing a variety of imprecise recombinants with sequence duplications. During the process of resolution these duplications may be lost via an *in cis* deletion process, or via further rounds of in trans recombination events.

In recent studies involving the next generation sequencing of whole cell RNA preparations from co-infected cells, and parallel CRE-REP assays using highly modified templates, we provide further evidence that recombination *per se* (the strand transfer event) is not influenced by either the RNA donor or acceptor sequence or its structure [13]. At face value this contradicts numerous studies of recombinant virus genomes which have implicated local RNA structure, sequence identity or particular nucleotides at the recombination junction [14–20]. However, progeny recombinant virus genomes in these studies are likely to have undergone one or many rounds of resolution and, in contrast to the initial recombinant viral RNA population in the cell cytoplasm, are – by definition – functionally intact and packaged into infectious virus.

It therefore remained a possibility that recombination and resolution are mechanistically dissimilar events, in which case sequence identity, composition or structure might influence one and not the other. To address this we used a well-characterised imprecise recombinant genome, #105B a type 1/3 poliovirus recombinant containing a 249 nt genome duplication [10], to investigate resolution independently of the initial template-switching event of recombination. We demonstrate that resolution of the virus population is driven by viral fitness and is enhanced in the presence of ribavirin, a mutagen that decreases RdRp fidelity and has previously been demonstrated to enhance recombination [10]. In addition, we characterised the resolving/resolved virus population following co-infection of uniquely barcoded imprecise recombinant viruses. We determined that resolution is, at least in part, a process requiring further template-switching/recombination events. The biphasic recombination process therefore appears to involve iterative strand-transfer events in which the primary determinant of selection is the fitness of the resulting virus genome.

## Results

### Rate of resolution is influenced by virus fitness

To better understand the resolution process, we investigated whether resolution was influenced by the sequence background of the imprecise recombinants studied to date. We have previously identified and subsequently constructed a cDNA for a viable intertypic (PV3/PV1) imprecise recombinant designated, #105B (GenBank MZ165620). The #105B cDNA has a 249 nt sequence insertion at the VP1/2A cleavage boundary consisting of duplicated VP1 and 2A sequences, as well as a fragment of firefly luciferase sequence derived from the sub-genomic replicon used in the CRE-REP assay in which #105B was generated (Fig 2A, upper panel) [10].

**Fig 2.**
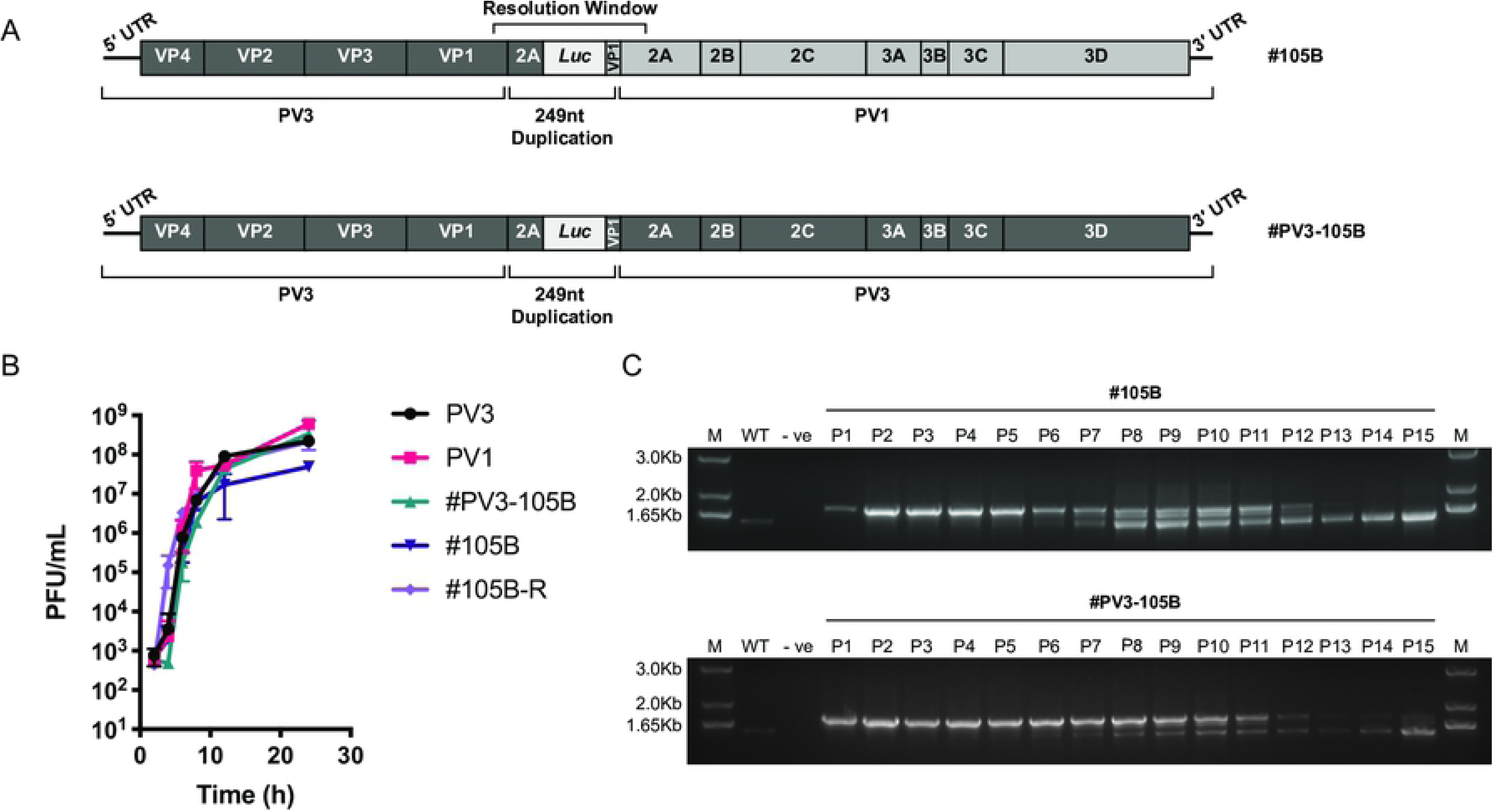
Phenotypic characterisation of intra- and intertypic imprecise recombinants. (A) Schematic representation of the genomes of imprecise recombinants #105B and #PV3-105B. Regions in dark grey are derived from PV3 and regions in light grey are derived from PV1. (B) Replication kinetics of imprecise clones. HeLa cells were infected with virus stock at multiplicity of infection of 5 and virus-containing supernatants harvested over 24 h. Virus titres were determined by plaque assay on HeLa cells. Error bars represent the standard deviation of 3 experiments. (C) Resolution of imprecise recombinant clones. Both #105B (upper panel) and #PV3-105B (lower panel) were serially passaged on HeLa cells for 15 passages. Viral RNA was extracted from supernatants and the potential region of recombination amplified by RT-PCR. The WT lane represents the 1470 nt product generated from a correct-genome length sample.

We reasoned that any *in cis* resolution events might be influenced by sequence identity even if the original strand transfer event had not been. We therefore generated an intratypic (PV3/PV3) version of #105B for comparison, designated #PV3-105B, containing an identical sequence duplication to that of #105B but in a sequence background of PV3 only (Fig 2A, lower panel; GenBank MZ165621). Analysis of the growth kinetics of #105B and #PV3-105B showed a small but significantly lower (P<0.05, t-test) average maximum titre of 4.9×10^7^ pfu/ml for #105B when compared to 2.2×10^8^ and 6×10^8^ pfu/ml for parental viruses PV3 and PV1 respectively (Fig 2B). In contrast, we observed no statistically significant loss in replication fitness of #PV3-105B, with a maximum virus titre of 3.3×10^8^ pfu/ml. This suggests that contrary to our original finding that imprecise recombinants have an associated fitness cost presumed due to the presence of sequence duplications, duplications alone may not have an observable effect on virus fitness.

We next tested whether these differences in replication kinetics influenced the rate of resolution of the two imprecise recombinant viruses. Duplicate samples of #105B and #PV3-105B were serially passaged 15 times (p1-15) on HeLa cells and the region spanning the imprecise recombination site was amplified by RT-PCR from cell culture supernatant at each passage (Fig 2C). Resolution of imprecise recombinants is reflected in the increased gel mobility of the PCR fragment spanning the recombination site, as deletions accumulate in the virus population. By p13, #105B had resolved to yield a single visible product representative of viruses with genomes of native ‘wild type’ length. In contrast, with #PV3-105B a minor product representing a greater-than-genome-length virus population was observable alongside resolving products up to, and including, p15. In both assays DNA products of intermediate size between unresolved and fully resolved products were visible. These were most prominent between p8 and p11 for #105B, and p9 and p11 for #PV3-105B (Fig 2C) and their gradual disappearance was suggestive of either an iterative resolution process or that they were ‘dead-end’ products subsequently out-competed in the virus population.

Analysis of the growth kinetics for the resolved #105B virus population (#105B-R) at p15 showed restoration of parental levels of virus replication, with a peak virus titre of 2.1×10^8^ pfu/ml (Fig 2B). This demonstrated that at the population level, resolution is in part driven by virus replication fitness, with fitter imprecise viruses, i.e., #PV3-105B, under less selective pressure. We would therefore expect fitter imprecise recombinants to persist in the population for longer than those with a replication defect.

### Resolution of a clonal recombinant virus generates a diverse virus population

We have previously reported apparently intermediate products in the resolution process [10] similar to the transient PCR products that appeared between p8 and p11 during the resolution of #105B and #PV3-105B (Figure 2C). To characterise these intermediate products in greater detail and so gain a better understanding of the overall virus population during resolution, we utilised Oxford Nanopore Technologies (ONT) high throughput sequence analysis of both #105B and #PV3-105B.

We selected individual or pooled passaged samples that would best represent the unresolved (p1) and intermediate resolved (#105B, p8-11; #PV3-105B, p10-13) populations. We also analysed #105B and #PV3-105B at p15. Furthermore, to investigate how a population that appears to be resolved may continue to evolve over time we additionally passaged #105B to p18 and included this population in the analysis.

An amplicon library was prepared according to Oxford Nanopore protocols for 1D analysis. The library consisted of uniquely barcoded PCR products, generated by amplifying a ∼1.74 Kb region spanning the resolution window (see Materials and Methods), and representing the unresolved, intermediate, and resolved virus populations (Table S1). The library was prepared such that those populations predicted to contain the most diversity in the range of recombinant sequences generated, had greater representation within the library.

Two independent libraries were generated, one for each set of passaged samples, and sequenced separately (reads available at SRA, BioProject PRJNA731157). For the analysis we opted to focus only on genomes that could yield viable infectious progeny virus, i.e., those in which any deletion still maintained the open reading frame. Any one deletion, however, could occur anywhere within the duplicated region of the resolution window spanning nts 3361-3720 (Fig 2A). Theoretically there were 14,209 possible deletions that fulfilled these criteria. These were created *in silico* and used as reference templates for alignment of the ONT reads (Fig S1; and see Materials and Methods). After optimising the alignment of individual reads to the template population, individual consensus sequences were generated representing the resolving or resolved genomes within the virus population (Fig S1).

As expected, the p1 populations for both #105B and #PV3-105B yielded single consensus sequences matching the input virus, supporting the validity of the analysis pipeline. In all other populations recombinant variants were designated as either fully resolved (i.e., the overall genome length was identical to that of the parental viruses due to the acquisition of a 249 nt deletion) or partially resolved (i.e., intermediate products in which some of the original 249 nt duplication was retained). Fully resolved, precise junctions for #105B were designated after the nucleotide position at which the recombinant genome sequence switched from matching the 5′ PV3 sequence to the 3′ PV1 sequence, with recombinants designated as the last nucleotide matching PV3 (e.g., PV3^3378^). However, due to sequence identity (81%) between PV3 and PV1 in this region there was often ambiguity as to the exact crossover point, as previously described for such precise junctions [10]. Resolution to a precise junction required the deletion of 249 nt, which, to remain viable, must include the 137 nt of luciferase sequence derived from the PV1 donor. The remaining 112 nt, encompassing duplicated regions of VP1 and 2A, could be deleted from either the PV1 donor-derived or PV3 acceptor-derived sequence (resolution window; Fig 2A). Consequently, a precise junction could theoretically occur at any one of these 112 nt positions. However, due to the level of sequence identity between PV1 and PV3 in the VP1/2A duplicated region only 23 precise junctions could be unambiguously defined. As fully resolved #PV3-105B is indistinguishable from parental PV3, precise junctions could not be defined. In comparison, recombinants of both #105B and #PV3-105B maintaining even a small sequence duplication could inevitably be determined more accurately. These partially resolved variants were designated by the number of nucleotides deleted from the initial 249 nt duplication (e.g., #105BΔ162).

Despite the high numbers of theoretical recombinants, analysis of the #105B populations, detected only seven partially resolved (Fig 3A) and eight fully resolved (Fig 3C) unique junctions (Fig S2; Table S2). Despite ∼32% of the total ONT reads being generated to the partially resolved #PV3-105B samples, only 3 unique junctions (#PV3-105BΔ231, -Δ225 and -Δ192) were present with sufficient frequency to generate representative consensus sequences (Fig 3B). Notably, a few junctions were conserved in both #105B- and #PV3-105B-derived resolving samples; #105BΔ225 and #105BΔ192 have identical deletion sizes and junction positions to #PV3-105BΔ225 and #PV3-105BΔ192. In addition, #105BΔ231 and #PV3-105BΔ231 contain an identically sized deletion, although the junctions are in different genome locations (Fig S2).

**Fig 3.**
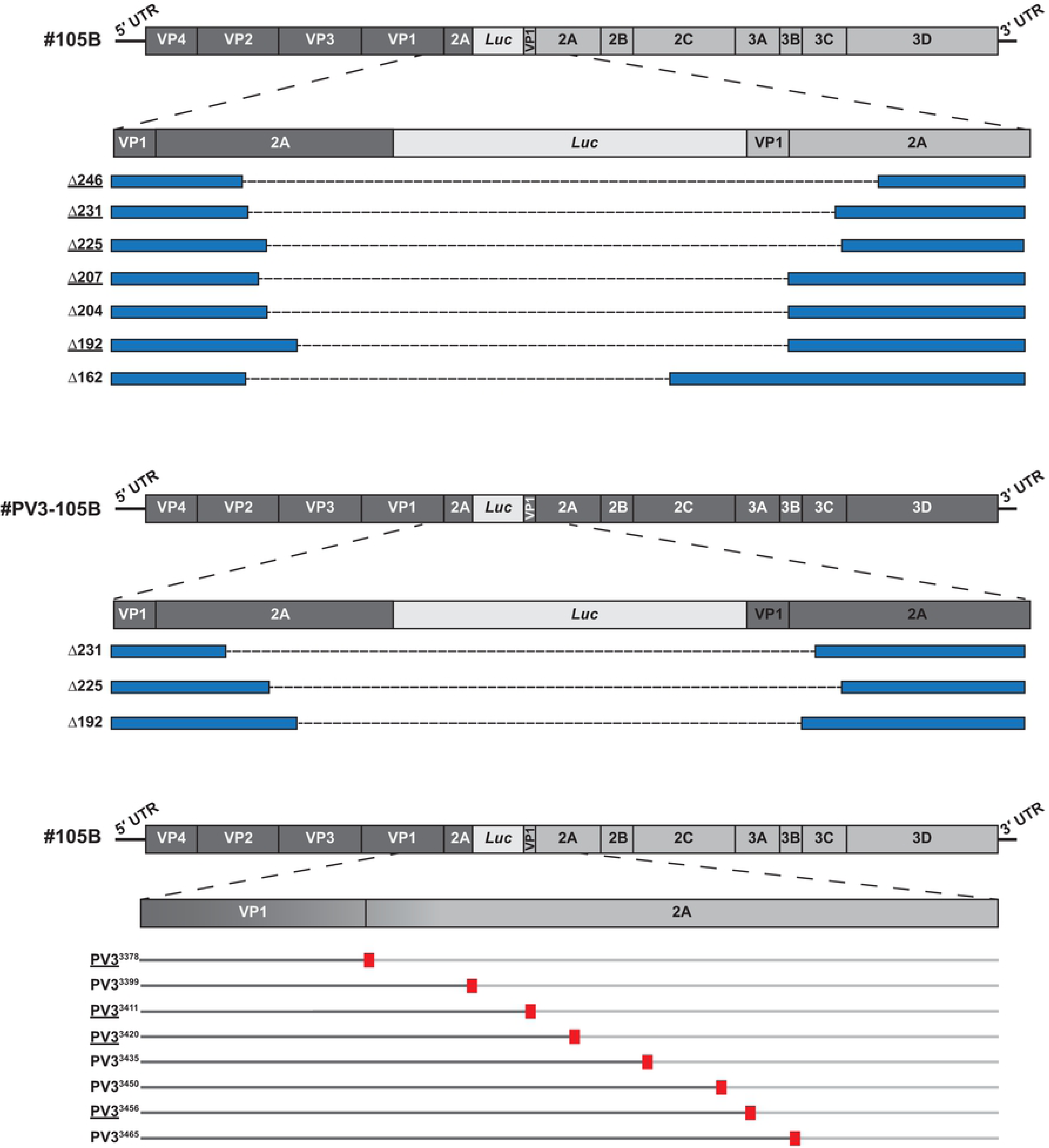
Nanopore analysis of #105B and #PV3-105B recombinant populations. The sequences of viable recombinant genomes were determined by consensus generation from Oxford Nanopore reads from all #105B and #PV3-105B passaging populations sequenced. Partially resolved genomes are shown for (A) #105B and (B) #PV3-105B. The original genome is shown at the top, where regions in dark grey are derived from PV3 and regions in light grey are derived from PV1, and an expanded view of the duplicated region shown below. Individual recombinants are denoted blue boxes with dashed lines representing the region of the duplication that has been lost. Numbers on the left-hand side represent the number of nt deleted from the 249 nt required to fully resolve the genome. Underlined recombinant names indicates recombinants detected in both replicate samples. (C) Resolved variants of #105B. Top schematic represents the genome of #105B with an expanded view of the VP1/2A region expected for a resolved recombinant. Resolved variants are represented by dual-coloured lines with sequence from PV3 in dark grey and sequence from PV1 in light grey. Red boxes denote the location of the resolved junction. Recombinant names, denoting the junction position, are given on the left. Underlined recombinant names indicates recombinants detected in both replicate samples.

Considerable overlap was observed in both the resolved and partially resolved genomes detected between the independently generated and analysed samples derived from #105B (Fig 4). Four of the fully resolved sequences (PV3^3378^, PV3^3411^, PV3^3420^ and PV3^3456^) and five of the partially resolved sequences (#105BΔ246, -Δ231, -Δ225, -Δ207 and -Δ192) were identified in both independent replicates. Unexpectedly, we identified the presence of several partially resolved variants at p15. Products representative of these variants were not visible during gel analysis (Fig 2C, upper panel); however, this is most likely a consequence of the limited sensitivity of the gel analysis technique. A subset of these partially resolved variants also continued to be present at p18, indicating that even seemingly resolved populations may be continually evolving alongside recombinant variants with non-wild type length genomes.

**Fig 4.**
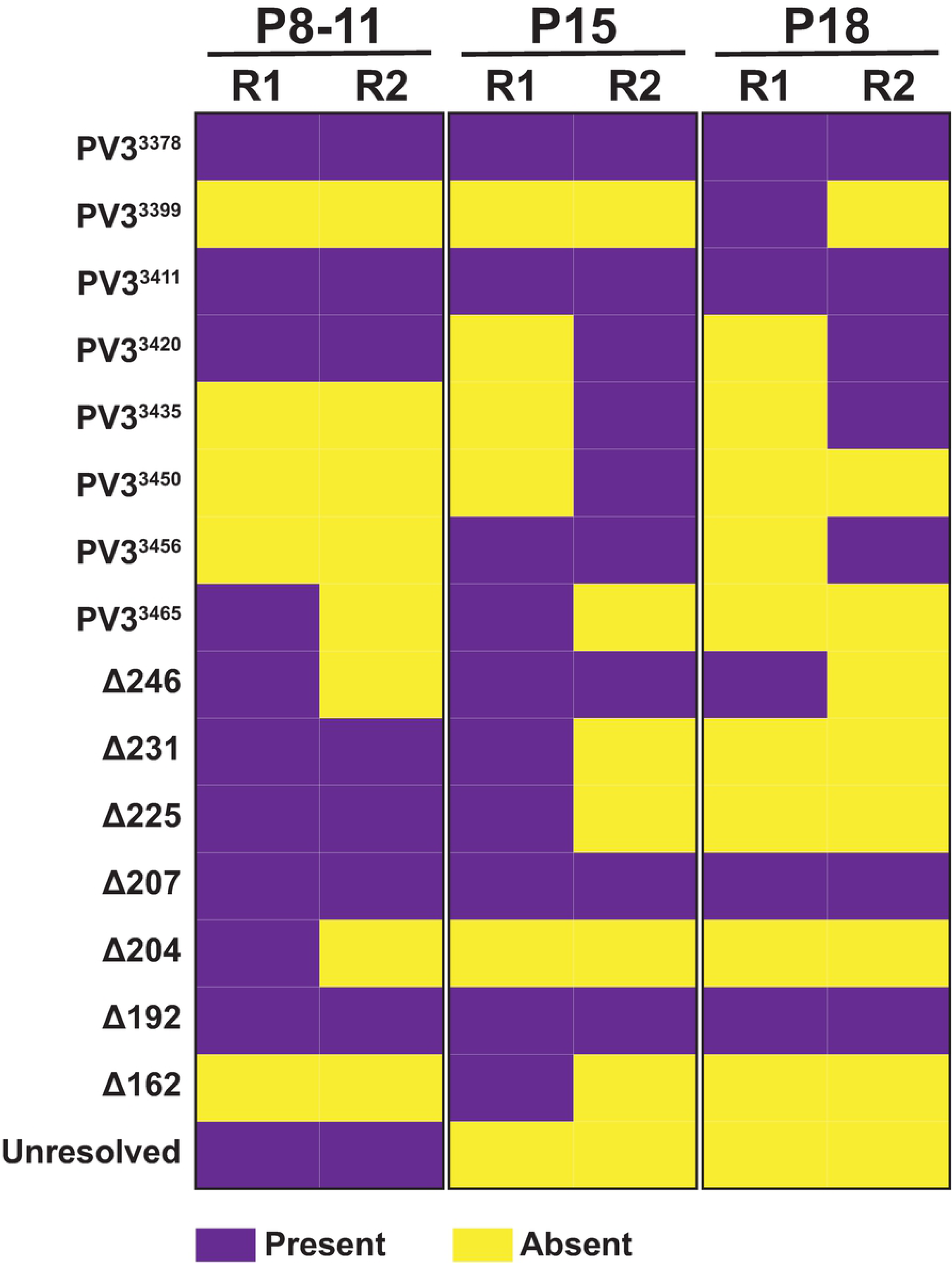
Comparison of recombinants identified between #105B nanopore replicates. Each recombinant variant identified from Oxford Nanopore analysis of the three #105B populations, p8-11, p15 and p18, are denoted as either present (purple) or absent (yellow) from passaging replicates (R) 1 and 2.

While strict selection criteria were applied to the data to minimize errors in recombinant junction or genome prediction caused by the high ONT error rate [21], these results suggest only a limited number of the theoretical viable variants were either generated or replicated to detectable levels during sequential virus passage. The similarities between partially resolved variants of #105B and #PV3-105B also suggested that specific recombinants were favoured regardless of the sequence of the backbone virus. As this was surprising, we used traditional Sanger sequencing to provide an independent analysis of the resolving population to determine that the ONT consensus sequences generated were accurate representations of the genomes extant in the virus population. Following PCR cloning, individual virus sequences were isolated from p9 and p18 of #105B, and p10 for #PV3-105B. Of the 20 clones analysed we were able to identify resolved variants PV3^3365^ and PV3^3399^, and partially resolved variants -Δ246, -Δ231, -Δ225 and -Δ192 from #105B, and partially resolved variant - Δ192 from #PV3-105B. Since these junctions were also all identified in the ONT analysis we concluded that consensus generation in the latter was representative of the underlying population of resolving recombinant molecules.

### Analysis of resolution by Illumina sequencing

The analysis of the resolving virus population by ONT showed it has significant promise in the study of recombination. For example, the long-read lengths enabled unresolved, partially resolved and fully resolved genomes to be detected in a single population. However, the high error rate inherent in ONT sequencing necessitated the use of strict criteria during pre-screening of primary data for consensus generation, including a threshold number of reads needed to form the consensus (see Materials & Methods). Inevitably this reduced sensitivity which we suspected resulted in missing less frequent resolving intermediates. In a recent study of recombination we had used paired-end Illumina sequencing to detect recombination junctions within total cell RNA. In this we were able to detect a much greater range of crossover junctions in the RNA population than were present in the resulting recombinant infectious virus population [13]. We therefore applied this method to investigate the process of resolution and to compare with results from the ONT analysis.

The imprecise recombinant #105B was passaged on HeLa cells until complete cytopathic effect (CPE) was observed for each cell monolayer, whereupon total RNA samples were harvested for subsequent RT-PCR and agarose gel analysis of the duplicated region as above. Under the conditions used in this assay, resolving genomes were apparent by p4 and resolution appeared near-complete by p10. After two further passages the virus populations from passages p2, p7 and p12 were selected for further analysis. In addition, a p0 sample – obtained following transfection of RNA – was analysed to determine the population diversity detectable prior to passaging. In contrast to the ONT analysis, which involved alignment to in-frame templates generated *in silico*, no constraints were placed on the resolving sequence junctions in the Illumina dataset which were identified using a bioinformatics pipeline based on ViReMa [13, 22, 23] (Fig S3; reads available at SRA, BioProject PRJNA731157).

The unpassaged p0 population, harvested 24 h post-transfection was not clonal, but instead contained a range of resolved precise deletions, together with additional genomes bearing imprecise insertions or deletions (Fig 5A). By p2, the total amount of variation within the sequenced population had markedly increased, with subsequent decreases at p7, and again at p12 (Fig 5A, B). Changes in population diversity followed a similar pattern when measured both as the number of previously undetected unique junctions identified at each passage (Fig 5B), and as a measure of Shannon Entropy (Fig S4). The level of #105B within the population was also reduced following each passage, as expected for the process of resolution (Fig 5B). A significant proportion of the junctions identified by VirReMa within the Illumina dataset were out-of-frame (∼30-50% in p0, p2 and p7), with the proportion dropping to ∼10% by p12 (Fig 5C).

**Fig 5.**
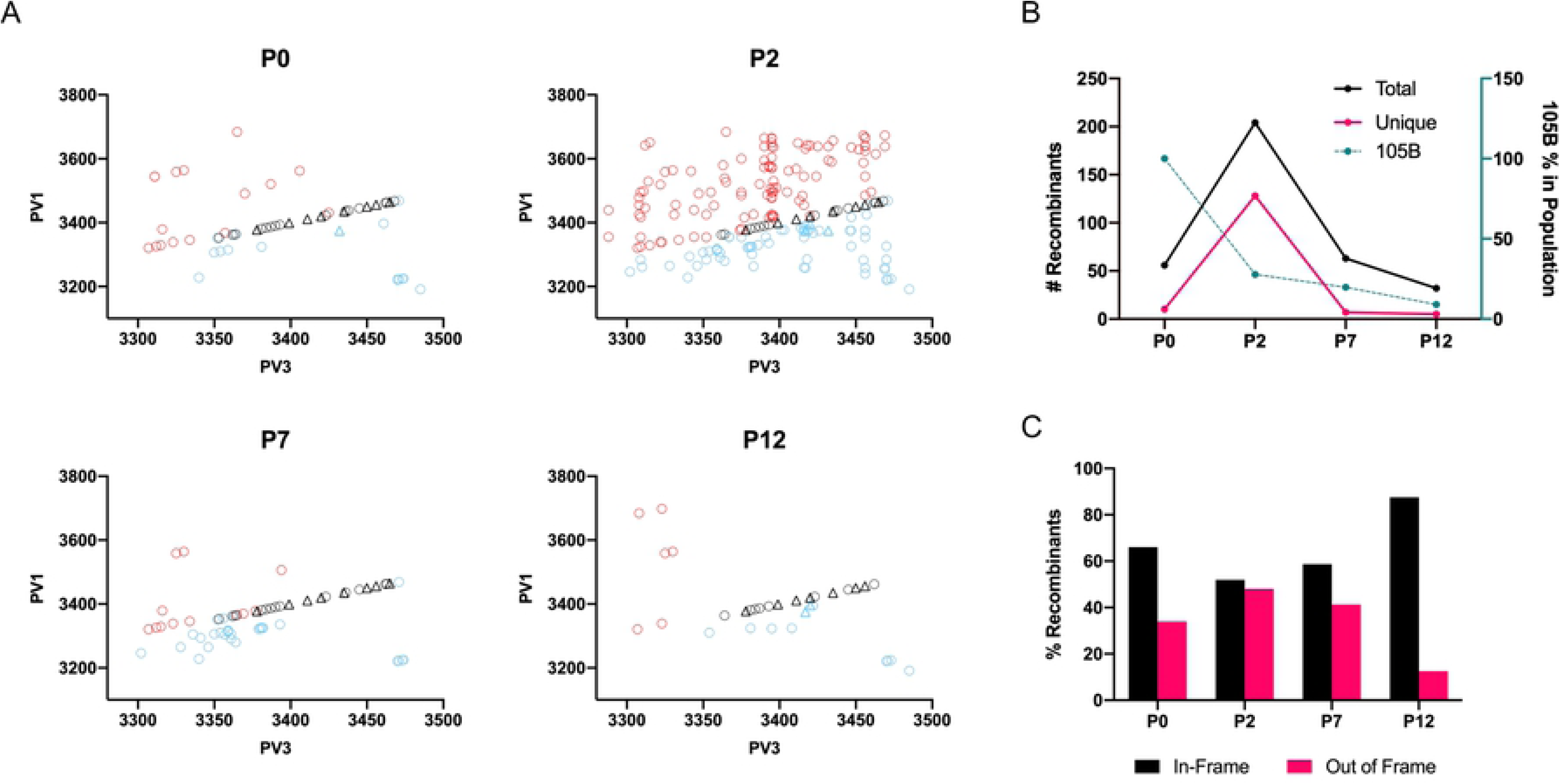
Visualisation of #105B Illumina recombinant type. (A) Recombinants detected via Illumina sequencing were plotted by type for p0, p2, p7, and p12; precise (open black circles), imprecise-insertion (open blue circles) or imprecise-deletion (open red circles). All recombinants detected with Oxford Nanopore as well as Illumina sequencing, irrespective of passage number, are indicated as precise (open black triangles) or imprecise-insertion (open blue triangles) recombinants. (B) Recombinant counts. The total number of independent junctions at each passage (left Y axis) was plotted against the passage number (x-axis) highlighting the level of diversity within the population at each passage. The proportion of #105B at each passage was plotted as a percentage of the total reads for #105B at p0 and after normalisation of all reads to p0 (right Y axis). (C) Comparison of the percentage of in-frame and out-of-frame recombinants present at each passage.

It was clear from the p0 data that, if resolution is a stepwise process, it can occur extremely rapidly as all of the 23 fully resolved variant genomes that can be unambiguously identified (as discussed above) could be detected within the 24 h period between RNA transfection and RNA extraction of the p0 sample. During subsequent passage the number of resolved variants detected fell from 23 at p0 to 15 at p12, presumably as the fittest variants came to dominate the population.

Overall, the Illumina sequence analysis of the RNA population during resolution revealed a comparable abundance, variety, and distribution of recombinant genomes as we identified during our study on recombination [13], indicating the same, or similar, mechanisms may underpin both processes.

### Ribavirin increases rate of resolution

RNA polymerase fidelity is a major determinant of recombination [10, 24], with error-prone RdRp variants generating recombinants at an increased frequency [25, 26]. One way to increase the error-rate of the native polymerase is to introduce sub-lethal concentrations of ribavirin, a well characterised guanosine analogue which induces base mismatches when incorporated into the viral genome and, at high levels, results in lethal mutagenesis by error catastrophe [27]. We have previously demonstrated that ribavirin increases the yield of recombinants in the CRE-REP assay by ∼300% [10]. If the process of resolution also involves additional rounds of recombination or is influenced by the error-rate of the RdRp, we predicted that the presence of ribavirin would also increase the rate of resolution.

We therefore serially passaged #105B, recovered following transfection with *in vitro* produced RNA, in the presence of a range of sub-lethal concentrations of ribavirin and monitored the resolving population by RT-PCR. There was a clear dose-response with increasing levels of ribavirin resulting in the more rapid appearance of a fully resolved product with increased mobility when analysed by agarose gel electrophoresis (Fig 6). In addition, the presence of the large unresolved parental #105B product – still apparent at passage 10 in the absence of ribavirin – was also influenced by increasing concentrations of ribavirin, being lost by p5 or p4 with 100 mM or 200 mM ribavirin respectively. These results indicate that the process of resolution is influenced by the error rate of the viral polymerase and is at least suggestive that further rounds of *in trans* recombination might be involved in resolution. To investigate the latter further we explored the generation of recombinant genomes during the resolution of uniquely barcoded variants of #105B.

**Fig 6.**
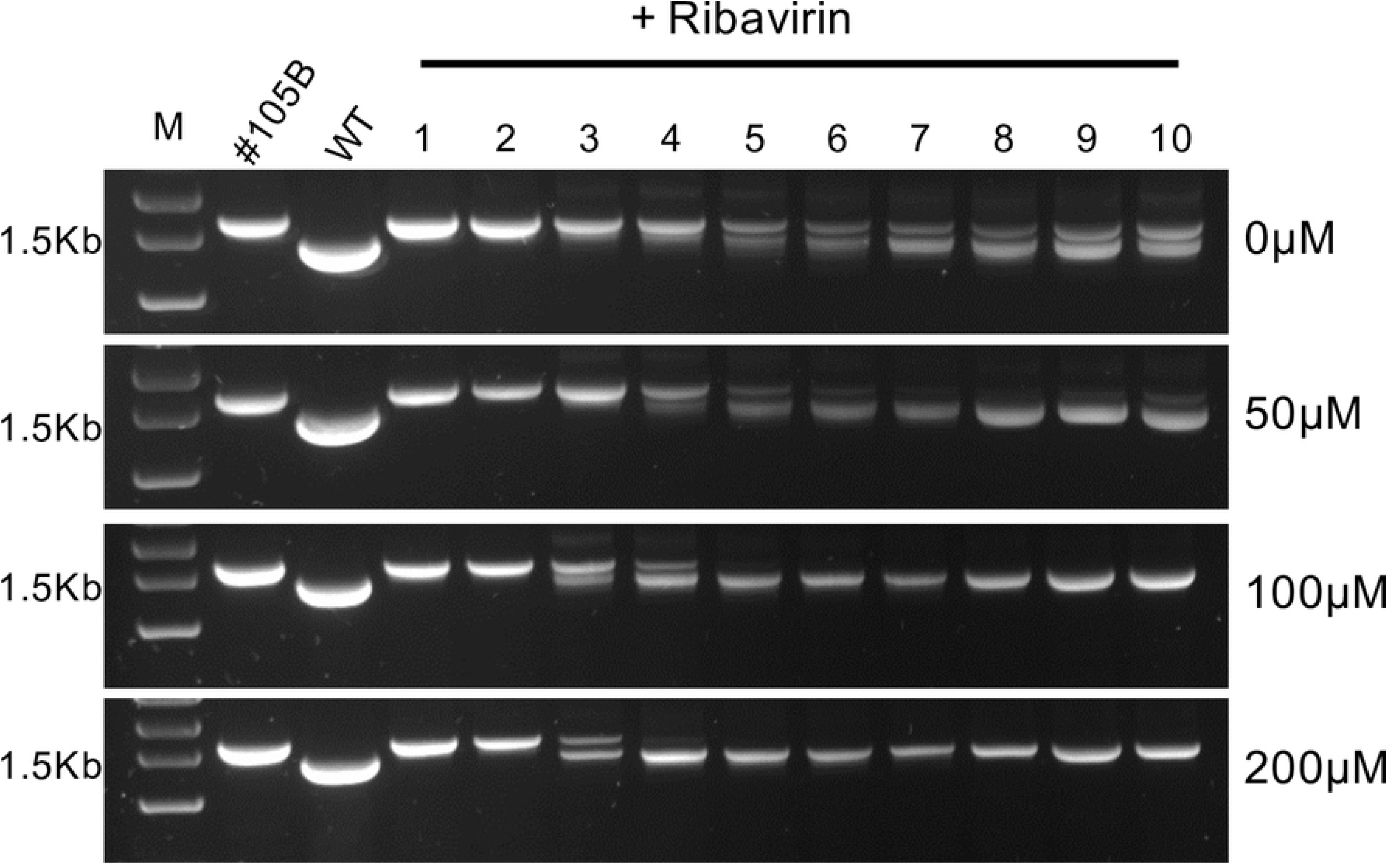
The rate of resolution of #105B increases in the presence of Ribavirin. #105B was serially passaged ten times on HeLa cells in the presence of 0, 50, 100 or 200 mM of ribavirin. Viral RNA was extracted, amplified by PCR and analysed by gel electrophoresis. WT = PV3, representing genome-length product. M = GeneRuler 1kb plus DNA markers (FisherScientific).

### Recombination is detected during resolution of barcoded viruses

The structure of the #105B genome means that all partially or fully resolved derivatives must effectively ‘lose’ sequences from within the resolution window. We therefore introduced modifications flanking this region that would allow the detection of resolving genomes that could only have arisen following a recombination event (Fig 2A; Fig 7A). Taking advantage of the degeneracy of the triplet amino acid code we synthesised a small panel of synonymous variants encompassing five-codons on each side of the resolution window. Individual viruses were recovered from engineered cDNAs using standard techniques and their growth kinetics compared in parallel to #105B. Two uniquely barcoded cDNAs designated #105B^T1^ and #105B^T2^ (Fig 7A), encoding viruses with replication kinetics indistinguishable from the parental #105B (Fig 7B) were chosen for further study.

**Fig 7.**
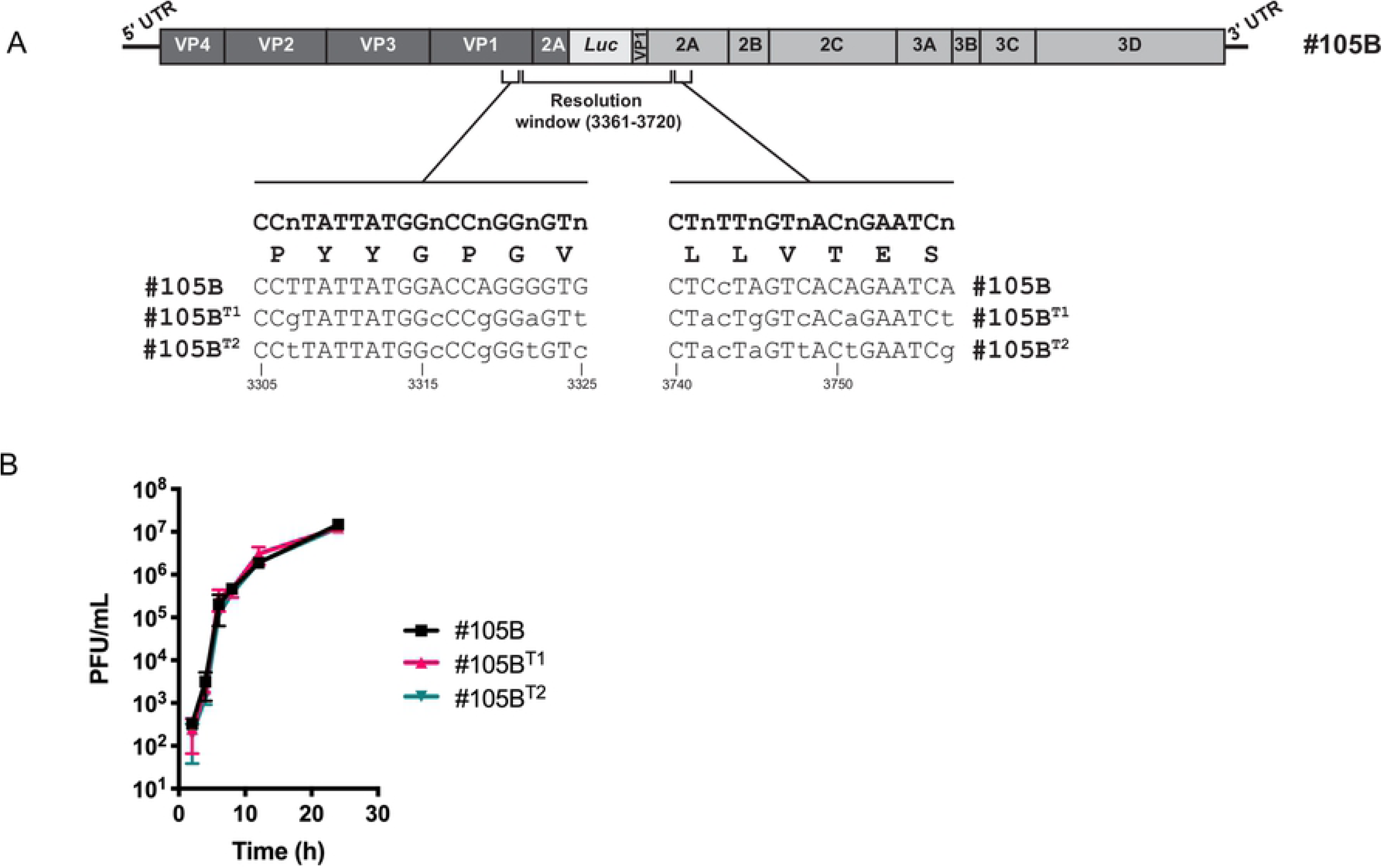
Characterisation of barcoded #105B. (A) Schematic representation of the sequences modified in #105B for the addition individual barcodes. (B) Replication kinetics of #105B and barcoded viruses.

HeLa cells were co-infected with #105B and either #105B^T1^ or #105B^T2^ at a 1:1 ratio of each virus and a final MOI of 10. Virus was harvested and serially passaged, with each passage titred to maintain the MOI of 10 in order to maximise the likelihood that each cell would be infected with at least one copy of each #105B variant. Resolution was monitored by RT-PCR amplification of the resolution window (and flanking barcodes), and 48 individually cloned PCR products from resolving, or resolved, populations were sequenced for each of the two co-infections (n=96).

Of the 96 sequences 89 consisted of a single clonal population, with the remaining 7 being mixed and excluded from further analysis. Of the 89 clonal sequences, 76 (85%) were unresolved (n=5), partially resolved (n=53) or completely resolved (n=18) with flanking barcode regions matching only one or other of the parental viruses. These neither supported nor refuted an involvement for recombination in the resolution process as the resolved genomes could have been generated by recombination between identical parental viruses or via an *in cis* deletion process. However, the remaining 13 sequences (15%), all of which were partially (n=11) or completely (n=2) resolved, contained mismatched sequences in the barcoded regions indicative of a recombination event. Four of these 13 sequences were biological duplicates, leaving a total of 9 unique resolving genomes that must have arisen from a recombination event (Fig 8; Table S3). Interestingly, 2 of the junctions identified – partially resolved -Δ207 and fully resolved PV3^3420^ – were identical to junctions previously observed when passaging #105B (compare Fig 3A and C to Fig 8), with -Δ207 also being detected from both barcoded co-infections. This observation was in agreement with the commonality of junctions observed between replicates of the #105B ONT analysis and again, implies that a limited number of resolving variants are detected from infectious virus supernatant, despite the abundance of variants at the RNA level [13]. Taken together these data confirm that resolution occurs, at least in part, via further rounds of template-switching events.

**Fig 8.**
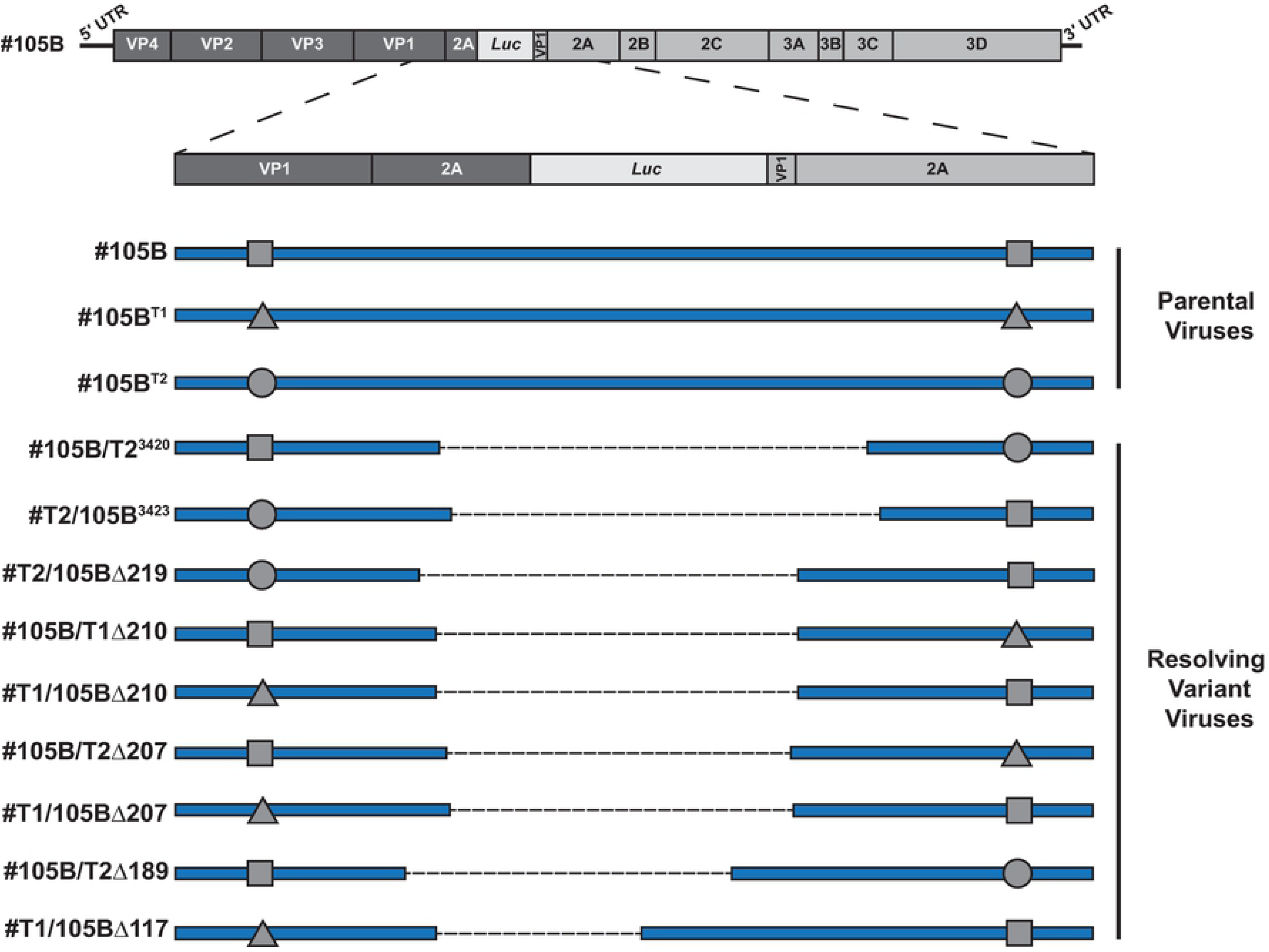
Analysis of barcoded #105B recombinants. Schematic representation of unique recombinant variants identified following co-infection of #105B with barcoded virus #105B^T1^ or #105B^T2^, and passage on HeLa cells until resolution was observed. Sequence names are given on the left with the top three, #105B, #105B^T1^ and #105B^T2^, representing parental constructs with modified sequences flanking the resolution window represented by a triangle (#105B^T1^) or a circle (#105B^T2^). Fully resolved or partially resolved sequences are shown with a dashed line representing the region that has been lost from the parental sequence, with appropriate shapes denoting the parental construct from which the 5′ and 3′ sequences were derived.

## Discussion

The evolution of RNA viruses with single-stranded, positive-sense genomes is, in large part, driven by the process of recombination. This process can overcome the effects of deleterious mutations on individual genomes, thus continuing to provide the population diversity required to prevent error catastrophe [28], while also resulting in the generation of novel viruses with potentially novel phenotypes [29, 30]. The mechanisms underpinning recombination have not been well characterised, but it appears to be a random and ubiquitous event that leads to considerable diversity at the genome level [13]. These initial polymerase template-switching events frequently result in the insertion or deletion of genomic sequence, referred to as imprecise junctions [10]. Although deletions - even if replication-competent - cannot give rise to viable viruses, imprecise insertions can be, and frequently are, replication competent and packaged into virus particles. We have previously demonstrated that viruses with imprecise insertions in the genome undergo a process – termed resolution – that leads to the loss of duplicated sequences and the creation of recombinant viruses with precise junctions and native or ‘wild type’ length genomes [10].

Recombination has previously been linked to the presence of RNA structures and/or regions of sequence identity [15-17, 19, 31-33], although there are a number of contradictory reports regarding the exact influence of such factors [18, 20, 34]. Our recent studies suggest that neither sequence identity nor RNA structure influence the polymerase template-switching event [13]. The majority of published studies on recombination have focused on analysing the end results of recombination i.e., viruses with wild type length genomes. If resolution is a distinct process from recombination, then it may be that the sequence and structure dependencies observed are a result of this second process and not the initial polymerase template-switching event. In the current study, we have therefore investigated this process of resolution in order to understand it better.

Throughout this study we have made use of a previously cloned imprecise intertypic type 3/1 poliovirus recombinant – #105B – containing a 249nt insertion spanning the P1/P2 junction boundary (Fig 2) [10]. To address whether sequence identity influenced the process of resolution we created an intratypic type 3/3 equivalent of #105B, designated #PV3-105B (Fig 2A), in which the duplicated viral sequences were all from poliovirus type 3. If sequence identity was a positive influence on resolution, we expected to see an increased rate of resolution for #PV3-105B when compared to #105B. Contrary to this expectation, we found that #PV3-105B resolved slower than #105B (Fig 2C). #PV3-105B also exhibited better replication kinetics compared to #105B (Fig 2B). As we have previously demonstrated that resolution selects for increasingly fit viruses [10] we conclude that the slower resolution rate of #PV3-105B is a consequence of initial increased fitness, with concomitant reduced selection pressure for better replicating variants. Although this confirmed that resolution is primarily a fitness-driven process, the differing replication rates meant we could not conclusively rule out a role for sequence identity in the process.

The gradual appearance of lower molecular weight products over the course of #105B and #PV3-105B passaging, leading to wild type length genome populations supports a sequential resolution process, as previously suggested [10], with some intermediate products being non-viable. To investigate these, we employed ONT sequencing of amplicons generated from virus supernatant from a passaged resolving virus population. Although the long read-length of nanopore technology allowed analysis of the entire amplicon, the inherent high error rate necessitated the development of an analysis pipeline that generated consensus sequences that matched one of the 14,209 in-frame - justified as all sequences were from supernatant virus - resolved genomes, which we generated *in silico* and used as templates (Fig S1).

We identified a relatively small number of recombinant variants in total, 15 for #105B and 4 for #PV3-105B (Fig 3). We confirmed by Sanger sequencing that our analysis pipeline was accurately detecting extant variants. We also observed that the ONT analysis was sensitive to the threshold (number of reads to generate a consensus) used to define a junction (Fig S1); reducing the threshold increased the range of junctions predicted, but introduced results potentially influenced by the template matched and the inherent high error rate of the technology. As the latter improves, the long ONT read lengths will likely offer the exciting potential for whole-genome recombination analysis.

One of the most interesting observations in the analysis of the #105B ONT data was the overlap observed in the variants isolated between different passaging replicates (Fig 4). The question arises as to whether these variants are (a) generated more readily or (b) have a fitness advantage, and therefore are amplified faster to detectable levels in the population. If resolution is a fitness driven process, then we would assume the latter. Resolved variant PV3^3378^ was detected in both replicates at all passages analysed (Fig 4), with the greatest number of consensus sequences, and therefore reads, generated, suggesting that this precise recombinant has a selective advantage. However, in direct competition assays with cloned recombinant PV3^3399^ – a resolved variant detected less frequently and at lower levels – no fitness advantage was observed for cloned PV3^3378^ (Fig S5). This implies that more complex population dynamics are at play than can be determined by the sensitivity of such an assay. The sequence identity between donor and acceptor means that the PV3^3378^ junction could occur anywhere between nucleotides 3366 and 3378. The consequence for the virus is that this may create a recombinant with a P1 region derived from PV3, and a P2/3 region from PV1. The intact and complete nature of the P1 and P2/3 ‘modules’ in such a recombinant may provide benefits in replication, packaging, or other aspects of the life cycle that lead to its competitive advantage and dominant prevalence in our *in vitro* assays.

Our ONT analysis of resolving viral genomes likely represented only a subset of the RNA variation present within the cell cytoplasm [13]. We therefore investigated the latter using Illumina next generation sequencing and observed significant levels of diversity within the RNA population (Fig 5A; Fig S3). We have previously demonstrated that the Illumina library preparation methods utilised here do not generate artefactual junctions that could confound the results [13]. We demonstrated a very good overlap between precise junctions independently identified using both ONT and Illumina technologies (Fig 5A).

As recently demonstrated for recombination [13], viruses undergoing resolution generate an extensive population of diverse molecules (Fig S3), the identity of which could arise from additional *in trans* polymerase-mediated template switching events. If that were the case, we would expect resolution to be influenced by factors known to influence recombination *per se*. Additionally we would also expect to be able to identify specific *in trans* crossover products with a suitable experimental system.

To test this hypothesis further we repeated the passaging of #105B in the presence of sub-lethal doses of ribavirin and analysed the resolving populations by gel electrophoresis (Fig 6). The link between reduced polymerase fidelity – resulting in increased mutation rates – and an increase in recombination is now well established and correlates with recombination being a driving force behind virus evolution and avoidance of error-catastrophe-based population decline. We observed an increase in resolution up to and including in the presence of 200 µM ribavirin, supporting our hypothesis that the mechanism underpinning resolution is further rounds of recombination. Notwithstanding this evidence, it remained possible that resolution occurs via a novel mechanism such as *in cis* deletion events, as increased mutations rates may also influence such a process.

By its very nature recombination requires two replicating genomes but, only when there are differences between those two genomes can recombination be identified. To discriminate between *in cis* deletion and *in trans* recombination events during the resolution of #105B we engineered unique ‘barcodes’ into two copies of #105B genome (Fig 7) and passaged each virus in the presence of wild type #105B. Analysis of the resolving virus populations identified recombination events between barcoded and non-barcoded #105B genomes (Fig 8) in 15% of the progeny analysed. Of these, the two junctions identified in the three variants #105B/T2^3420^, #105B/T2Δ207 and #T1/105BΔ207, had been observed previously (Fig 3A, C) supporting our theory that a select few recombinants with a replication advantage are likely to colonize a host.

Presuming recombination occurs during negative strand synthesis [4] the 3′ genome sequence can be regarded as being derived from the donor template, with the 5′ sequence derived from the acceptor. By its nature the CRE-REP assay could only generate viable genomes with a 5′ PV3 and 3′ PV1, and therefore it was not possible to establish if there was an inherent bias for templates to act as a donor or an acceptor. In the analysis of the barcoded recombinants we observed templates acting as both donor and acceptor based on the orientation of the 5′ and 3′ barcode sequences, with 5 examples of #105B as the donor and 4 examples of #105B as the acceptor, thus demonstrating, albeit with a limited number of samples, no obvious bias in the directionality of recombination as we have previously indicated [13].

It is clear, though perhaps unsurprising, that the resolution process can and does involve further rounds of recombination that necessitates an *in trans* strand transfer event. An iterative recombination process also explains the presence of out-of-frame recombinants continually detected throughout passaging of #105B (Fig 5C). As we have previously shown that recombination is a ubiquitous and apparently random process [13] out-of-frame junctions are possible when template-switching events occur at either position 1 or 2 of any individual codon and will be generated anew with each passage. However, as out-of-frame genomes will not go on to produce infectious progeny virions, and as shown in this study (Fig 5A), the relative percentage in the population will eventually be surpassed by the fitter, and replication competent, in-frame genomes.

While the evidence highlights a role for *in trans* template-switching events, it remains unclear as to whether *in cis* events are also driving the evolution of viruses with imprecise genomes. It is perhaps more appropriate to consider the process as the dissociation of the polymerase from the RNA template, and whether subsequent re-association occurs on the same (*in cis*), or different (*in trans*) templates. Using experiments with molecular tweezers enterovirus polymerases have been shown to pause and subsequently undergo back-tracking on the RNA template; in doing so producing a recombinogenic 3′-end, which can then be involved in template-switching – *in trans* – or copy-back type – *in cis* – recombination events [35] [36]. Such a mechanism would suggest that resolution occurs both *in cis* and *in trans* and perhaps the limited number of recombinants (15%), conclusively generated via template-switching events (Fig 8), is a consequence of the two interconnected processes at play. Such a mechanism would correlate with the idea that poliovirus polymerases form lattice structures within the replication complex [37]. Following dissociation, if the nascent RNA is able to freely move between polymerase molecules within the lattice, then recombination via template-switching may simply be a result of RNA movement towards the active site of a different polymerase molecule, while *in cis* recombination is due to less RNA movement and therefore re-association on the same RNA. Investigation of recombination in the context of higher order polymerase structures would offer valuable insights into our further understanding of this fundamental process.

## Materials and Methods

### Virus and cell culture

HeLa cells were maintained at 37°C, 5% CO_2_ in Dulbecco’s Modified Eagle Medium (DMEM; Sigma) supplemented with 10% heat-inactivated FBS (FBS-DMEM; Life Technologies). Poliovirus type 1 (Mahoney), type 3 (Leon) and recombinant pT7- #105B were recovered following transfection of an *in vitro* transcribed RNA from full-length cDNA and virus stocks quantified by plaque assay on HeLa cells. Growth kinetics of viruses were determined following infection of HeLa cells at a multiplicity of infection (MOI) of 5 in serum-free media (SF-DMEM). Unabsorbed virus was removed by washing with PBS and cells incubated in fresh FBS-DMEM at 37°C, 5% CO_2_. Supernatants containing virus were harvested at various time points post-infection and quantified by plaque assay on fresh monolayers of HeLa cells. Unless otherwise stated, for serial passaging HeLa cells were initially infected with virus at MOI 1, while for subsequent passages harvested supernatant was diluted 1∶4 with SF-DMEM prior to infection of fresh monolayers of HeLa cells.

### in vitro RNA transcription and transfection

PV1 and #105B variant plasmids were linearized with *Apa*I, and PV3 linearized with *Sal*I, and RNA transcribed using a HiScribe T7 High Yield RNA Synthesis kit (NEB) following the manufacturers’ protocol. RNA transcripts were DNaseI (NEB) treated to remove residual template DNA and column purified using a GeneJET RNA Purification kit (ThermoFisher) prior to spectrophotometric quantification. For virus stocks, 1 µg of RNA was transfected into 80% confluent HeLa cell monolayers using Lipofectamine 2000 (Life Technologies) in a 3:1 Lipofectamine 2000:RNA ratio as per manufacturers’ protocol. Virus supernatant harvested 24 h post-transfection and quantified by plaque assay on HeLa cells.

### Synthesis and infection of barcoded #105B

The region of #105B in which resolution could occur to yield a precise genome (resolution window) is located from nt 3361 to nt 3720 of #105B. Five amino acids, with maximum coding redundancy, were selected flanking either side of this region: 5′-PV3_VP1 Pro277, Gly280, Pro281, Gly282, Val283 and, 3′-PV1_2A Leu39, Leu40, Val41, Thr42, Ser44. A geneblock library was synthesised (IDT) with an n at the 3^rd^ base position of each amino acid. Additionally, Leu40 was altered from TTA > CTn to ensure maximum redundancy without altering the amino acid. Individual DNA fragments were used to replace the equivalent WT sequence in pT7-#105B using HiFi Builder cloning technology (NEB). Colonies were screened and sequenced and two plasmids containing unique and divergent sequences were selected as pT7-#105B^T1^ and pT7-#105B^T2^.

HeLa cells were seeded at 5 x 10^5^ per well in 6-well plates 18 h prior to assay. Cells were co-infected with virus inoculum at a final MOI 10 for 30 min at 37°C. Inoculum was removed and cells washed once with PBS. Fresh media was added and cells incubated for a further 8 h at 37°C, 5% CO_2_. Virus-containing supernatant was harvested and quantified by plaque assay on fresh HeLa cells prior to subsequent passage at MOI 10. RNA was extracted from viral supernatants at passages 7 and 9 and the resolution window amplified by RT-PCR as described below. PCR products were cloned using the pGEM-T Easy System (Promega) and sequence determined by Sanger sequencing (Eurofins Genomics, Germany) with primer VP1(F): 5′- GGTGTATTGGCAGTTCGTGTTG.

### RNA extraction and RT-PCR

Viral RNA was isolated from clarified cell culture supernatants using an Omega Total RNA Isolation Kit 1 (Omega BioTek, VWR) and reverse transcribed at 42°C using an oligo dT primer and SuperScript II reverse transcriptase (Life Technologies) as per manufacturers’ protocol. The resolution window was amplified using standard Taq polymerase (NEB) with an initial denaturing at 95°C for 30 sec, followed by 30 cycles of 95°C for 30 sec, 50°C for 30 sec and 68°C for 90 sec, and a final extension at 68°C for 5 min with primers PV3(F): 5′-GCAAACATCTTCCAACCCGTCC and PV1(R): 5′- TTGCTCTTGAACTGTATGTAGTTG, or PV3-IS(F): 5′- CCCAAACACGTACGTGTCTG and PV1-IS(R): 5′-TGTATCCGTCGAAGTGTGATGG for Illumina sequencing. PCR products were separated by 1% agarose gel electrophoresis and sequence determined by Sanger sequencing (Eurofins Genomics, Germany). Sequence analysis was completed using SnapGene, Version 4.0.8. Illumina sequencing was carried out and analysed as previously described.

### Ribavirin Treatment

HeLa cells were seeded at 2 x 10^5^ per well in 12-well plates and infected with #105B at MOI 1 for 30 min. Cells were washed with PBS and fresh media added to all wells. Ribavirin was added to a final concentration of 0, 50, 100 or 200 mM and cells incubated for 24 h. Virus was subsequently blind passaged by infecting fresh monolayers of HeLa cells with a 1:4 dilution of supernatant from the previous passage, with Ribavirin added as above. At each passage, RNA was extracted from viral supernatants and the resolution window amplified by RT-PCR as described above.

### Next generation sequencing library preparation and sequencing

Viral RNA was extracted from serially passaged #105B and #PV3-105B virus supernatants and reverse transcribed as described above. For Illumina sequencing, samples were prepared and reads analysed as described in [13].

For Oxford Nanopore Technologies (ONT) analysis, primers for the initial PCR amplification of the resolution window, were adapted to include the Oxford Nanopore adaptor sequence and reactions cycled as described above. PCR products were purified using a Wizard® SV Gel and PCR Clean-up System (Promega). Samples were barcoded using Oxford Nanopore PCR barcodes (EXP-PBC001) as per manufacturer’s protocol. PCR products were purified as above and quantified on a NanoDrop ND1000. Amplicons were pooled in desired ratios (Table S1) with a total of 1 µg of amplicons used to create a DNA library. Library construction was carried out as per Oxford Nanopore protocol SQK-LSK109. The library was run on a FLO-MIN106 flow cell primed as per protocol SQK-LSK109 with reads collected over an 8 h period.

### Oxford Nanopore Technologies sequence data processing

Raw fast5 files were basecalled and de-multiplexed using ONT Albacore Pipeline software (v 1.2.4). Passed reads were converted to FASTA format using the extract command in Nanopolish (v 0.6.1) [38].

A variants database was created of 14,209 potential in-frame, resolved/resolving sequences within the resolution window. ONT reads were screened against the variants database using Blastn. Reads with <90% sequence identity to a potential variant, and that were +/-10% of the min/max possible length of the PCR product, were removed. Remaining reads were extracted and ordered based on each potential variant and all the ONT reads which matched to it. Aligned reads were processed using Nanopolish to generate consensus sequences against best matched potential variants. A cut-off limit was applied, where only template matches which contained >100 ONT reads were processed to generate a consensus sequence. Using Nanopolish, all generated consensus sequences were verified as in-frame and confirmed as matching the best potential variant template. A recombinant was defined as true if ≥10 identical consensus sequences were generated. Scripts available on request.

## Acknowledgements

The authors would like to express their gratitude to Professor Nicola Stonehouse, Professor Mark Harris, Dr Lee Sherry, and members of the Stonehouse group for providing facilities and assistance at the University of Leeds for K.B to complete some of the experimental data presented in this paper.

